# The impacts of grazing pressure by large herbivores on short-tailed field vole (*Microtus agrestis*) population cycles: results from a long-term upland experiment

**DOI:** 10.64898/2026.01.08.698433

**Authors:** Matt Dopson, Nick Parker, Laura Wadkin, Andrew Baggaley, Robin J Pakeman, Stuart W Smith, Darren M Evans

## Abstract

Understanding how large herbivores influence cyclic small-mammal populations is essential for predicting community dynamics in grazed ecosystems. Using a 23-year upland grazing experiment in Scotland, United Kingdom, we test how varying livestock grazing pressures affect population cycles of the short-tailed field vole *Microtus agrestis*, a keystone herbivore and prey species of birds of conservation concern.

We combined long-term vole sign indices (VSI), vegetation surveys, climate data and used wavelet analyses and mixed-effects models to quantify changes in VSI (a proxy for abundance), cycle amplitude and periodicity across four grazing treatments and three spatial blocks.

Vole abundance and cycle amplitude declined with increasing grazing pressure, yet the frequency of vole population oscillations remained strikingly consistent across all treatments, with a dominant ~3.5-year cycle persisting throughout most of the study. Ungrazed plots supported the highest mean VSI and the largest amplitude oscillations. We found a pronounced, spatially-restricted cycle collapse occurred between 2009–2016 in one experimental block. This breakdown coincided with unusually cold winters and lower vegetation density, suggesting that local microclimate and habitat structure interact to destabilise vole cycles. Vegetation density (but not height) predicted VSI in the more intensively grazed treatments, indicating that grazing-driven changes in habitat structure indirectly mediate vole dynamics.

Our findings demonstrate that large herbivores can substantially modify vole population dynamics through both direct and indirect pathways, altering cycle amplitude, habitat suitability and resilience to climatic perturbations. These results highlight the need to consider interactions between grazing, vegetation structure and climate variability when predicting small-mammal population dynamics and their cascading ecological effects.

## 1 INTRODUCTION

The *Arvicolinae* subfamily of rodents is one of the most abundant groups of mammals in the world. With species including voles, lemmings and muskrats, their range spans the northern hemisphere (Wilson & Reeder, 2005). *Arvicolinae* rodents act as a keystone species in many northern ecosystems, as they are an important food source for a wide range of predators (Tamarin, 1985). In the UK, the short-tailed field vole (*Microtus agrestis*) is the most common mammal, with some estimates putting their numbers near 80 million individuals (Harris et al., 1995). This immense population size means that they have long been recognised as having a significant influence on the ecosystems they are part of (Summerhayes, 1941; Sirotnak & Huntly, 2000). In particular, they are the main food source of predators such as the least weasel (*Mustela nivalis*) and common kestrel (*Falco tinnunculus*), as well as being a large part of the diet in many other raptors and terrestrial carnivores (Dyczkowski & Yalden, 1998; Tapper, 1979). This includes being an important part of the breeding success in birds of concern such as the hen harrier (*Circus cyaneus*) (Evans et al., 2006). However, this is not their only role in the ecosystem - voles are often described as keystone species due to their role as ecosystem engineers (Evans et al., 2006), playing a key role in seed dispersal (Fischer et al., 2018; Sullivan & Sullivan, 2021), seedling and sapling mortality (Gill & Marks, 1991; De Steven, 1991; Mittelbach & Gross, 1984; Hambäck et al., 1998) and the creation of mosaic microhabitats (Questad & Foster, 2007; Howe et al., 2006).

Like many *Arvicolinae* rodents, short-tailed field voles are known for their extreme population cycles, which see population numbers increase up to tenfold and drop back down again in just 3–4 years (Krebs, 2013). These cycles were famously recorded by Elton in 1924, where during peak years he described “plagues” of voles decimating agricultural land (Elton, 1924). Although widespread crop destruction by voles is now less common, it still occurs in modern agriculture (Jareño et al., 2015; Jacob et al., 2014).

There has been extensive research into the mechanisms that drive vole population cycles, with numerous studies in particular in regions such as North America, Scandinavia and Japan (Krebs, 2013; Oli, 2019); however, most of these studies focus on tundra biomes, where regular snow cover is a key factor. Particularly in Fennoscandia, research increasingly supports the view that specialist and generalist predators are the main drivers of vole population cycles (Turchin & Hanski, 1997; Hanski et al., 1991, 1993; Hanski & Henttonen, 2002; Hanski et al., 2001). In more temperate regions, such as the UK and central/southern Europe, studies, including Lambin et al. (2006), have shown that the patterns observed in more northern experiments do not apply. Through a weasel removal experiment, Graham & Lambin (2002) demonstrated that the impact of specialist weasel predation on field-vole survival was not necessary to drive the cyclic pattern of field-vole populations in Kielder Forest, United Kingdom. Moreover, Lambin et al. (2006) argued that it is possible that small rodent population cycles could have different drivers in different regions. Indeed, the key drivers of vole cycles in the UK are still not fully understood (Lambin et al., 2025).

It is well known that livestock grazers such as domestic sheep and cattle, alter vegetation structure, including height, density, biomass and community composition, which affects how small mammals use the habitat (Long et al., 2017; Bakker, 2003). Large grazers have been shown to reduce small mammal abundance (Parsons et al., 2013). The Jarman– Bell principle proposes that as herbivore body size increases, the quality of their food intake tends to decrease; however, this lower quality is offset by the greater quantity of food consumed (Jarman, 1968; Bell, 1971; Geist, 1974). This principle suggests that voles require higher quality, more nutrient rich food than the livestock grazers they often share land with. Although it is clear that there is an interaction, between small and large herbivores, the mechanics behind such interaction are not fully understood.

The Glen Finglas grazing experiment, located in the Loch Lomond and The Trossachs National Park, Scotland, is a long-term ecological study investigating how different livestock grazing regimes influence upland biodiversity. Established in 2002, the experiment consists of a gradient of grazing intensities (and a mixture of sheep and cattle) from ungrazed to commercially grazed areas. Research to date, based on bi-annual surveys of vole sign indices (VSI, Evans et al. (2006)) has revealed significant relationships between vole populations, grazing intensity, and predator interactions, including those with red foxes (*Vulpes vulpes*) (Evans et al., 2015; Villar et al., 2014, 2013). However, these early studies have largely relied on short-term datasets and were unable to consider the impacts of large herbivores on vole cyclicity. In the present study, and with the benefit of over two decades of data, we examine how the impacts of livestock grazing pressure by large herbivores influence vole population cycles and their broader ecological dynamics. We explore the following hypotheses:

A. Livestock grazing pressure will significantly affect mean vole cycle amplitude and mean vole population. In the absence of large herbivores, and based on short-term observations (Evans et al., 2006), we predict larger amplitude cycles and higher mean vole sign indices (VSI);
B. At the landscape scale, there will be little variation in VSI across the experiment design as they have largely congruent vegetation structure and composition, and experience the same weather conditions;
C. Changes in the vegetation community as a result of livestock grazing will have a significant effect on VSI. Due to dietary overlap between sheep and voles, we expect higher levels of competition in intensively sheep grazed areas compared to extensively sheep grazed or sheep and cattle grazed areas;
D. Finally, it is possible that vole population dynamics at Glen Finglas have been affected by the widespread vole cycle dampening observed in many similar experiments across the northern hemisphere in the last 20-30 years (Cornulier et al., 2013; Hörnfeldt, 2004; Hörnfeldt et al., 2005; Ecke et al., 2017). We examine this potential across the long-term dataset and predict, if present, that effects will be more acute in intensively grazed areas.

## 2 METHODS

### 2.1 Experimental Set-Up

The ongoing experiment is being carried out at Glen Finglas in the Loch Lomond and The Trossachs National Park in Scotland, United Kingdom (56°16’N, 4°24’W), see Fig 1 (left). The Glen Finglas estate, now covering 4875 hectares, has historically been grazed predominantly by sheep, alongside occasional cattle and deer grazing. The estate is now managed by the Woodland Trust, whose primary management objective for the Glen Finglas estate is nature recovery alongside extensive tree planting; including, incorporation into their largest woodland regeneration project - the Great Trossach Forest (Woodland Trust, 2025). The primary habitat of Glen Finglas is upland acid grassland. This rough grassland is dominated mainly by *Molinia caerulea–Potentilla erecta* mire, *Juncus effusus/acutiflorus–Galium palustre* rush pasture and *Festuca oviana– Agrostis capillaris–Galium saxatile* grassland communities (Pakeman et al., 2019). Naturally, Glen Finglas is hilly, hence all of the experimental plots are located on sloped ground at varying elevations from 200-500m above sea level (Evans et al., 2015). As this is an upland environment, coupled with the lack of nearby trees, all sites are exposed to wind and rain year round. The different blocks are also positioned at slightly different heights with different aspects. The main focus of the experiment is understanding how varying grazing treatments affect local flora and fauna.

**FIGURE 1.**
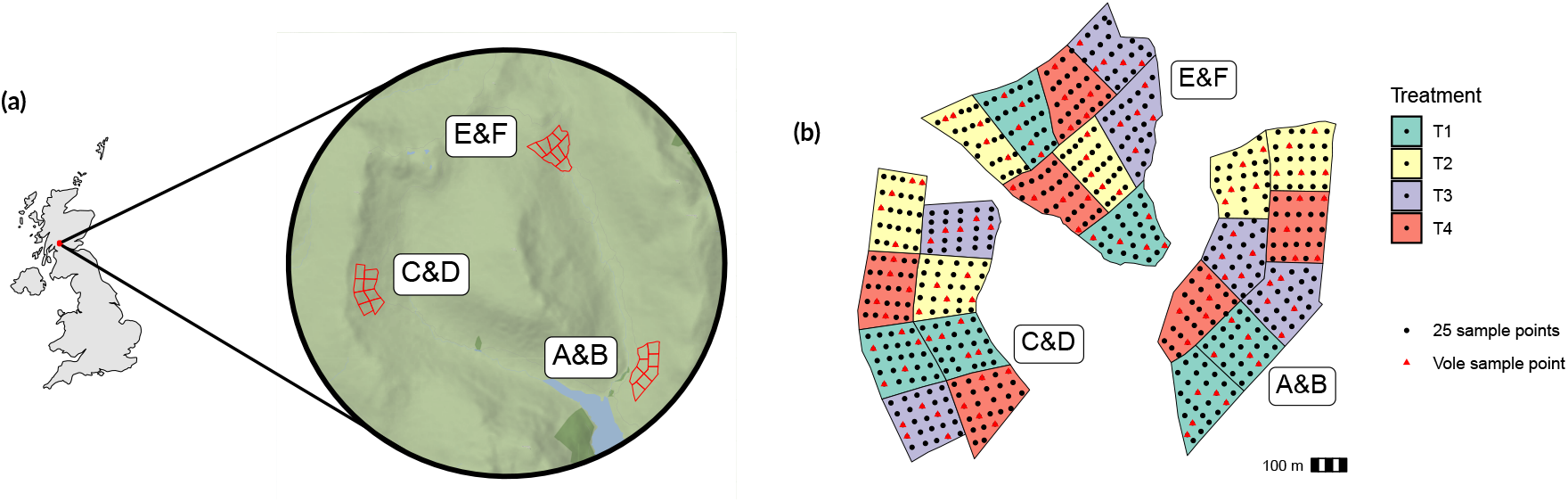
**(a)** Map of plots (shown as the individual red polygons) in Glen Finglas, shown within the UK (created using ggmap (Kahle & Wickham, 2013)). All three blocks, labelled as A&B, C&D, E&F, are situated on hillsides, with block C&D being on the steepest hill. **(b)** Spatial configuration of sample points within plots in Glen Finglas. Please note that the blocks (A&B, C&D and E&F) have been moved closer together to make the figure clearer. The points (black circles or red triangles) are the 25 sample points in each plot, with red triangles indicating the five points per plot where voles are sampled.

A replicated, randomized-block experiment consisting of six replicates of four treatments was initiated in 2003, with baseline data collected in 2002. Three blocks of land were chosen as the site of the experiment. These will be referred to as block A&B, block C&D and block E&F, shown in Fig. 1 (left). Each block contains two replicates of the experiment, e.g., block A&B contains replicates A and B. Each replicate contains four grazing treatments: T1, T2, T3 and T4. These range from conventional highland sheep-grazing levels (T1) through to completely ungrazed (T4). A summary of the different treatments is shown in Table 1. Accounting for plot size, this corresponds to nine ewes per plot in T1, three ewes per plot in T2, two ewes and four weeks of two cows and two un-weaned calves per plot in T3 (equivalent off-take to T2), and no livestock in T4. Each individual experimental plot is a subset of a block. The plots are contained in their own individual 3.3ha fenced field and contain only one treatment type. The geographic layout of the plots is shown in Fig. 1 (left), where each individual coloured polygon represents one experimental plot. Each plot contains 25 evenly spaced vegetation and invertebrate sampling points approximately 40m apart, with points at the edge approximately 20m from the fences. Within each plot, five points were randomly selected for vole surveys, and these same points are surveyed every year (see coloured points in Fig 1) (right).

**TABLE 1.** Summary of the four grazing treatments at the Glen Finglas long-term upland grazing experiment.

| Experimental treatment | Grazer stocking levels |
| --- | --- |
| T1 | 2.72 ewes $\text{ha}^{-1}$ (conventional stocking rate) |
| T2 | 0.91 ewes $\text{ha}^{-1}$ (one third conventional rate) |
| T3 | 0.61 ewes $\text{ha}^{-1}$ plus, for four weeks of each year, two cows with two un-weaned calves (one third conventional rate off-take) |
| T4 | 0 ewes $\text{ha}^{-1}$ (ungrazed) |

Twice per year, in April and October, a vole survey was conducted (except for April 2005, October 2017, April and October 2020, October 2022 and April 2023 when surveys were not feasible for logistical reasons). During the survey, five quadrats (25cm x 25cm) are randomly thrown and searched for vole sign indices (VSI) at each of the vole survey points. VSI includes searching for signs of vole runways, fresh vole droppings and fresh vegetation clippings caused by vole foraging (Evans et al., 2006, 2015). VSI are used here as a proxy for vole abundance as VSI have generally been shown to be linearly related to actual vole densities based on snap-trapping methods and are widely used to estimate vole abundance (Lambin et al., 2000; Wheeler, 2008; Jareno et al., 2014; Hansson, 1979; Redpath et al., 1995). VSI hereafter refers to vole droppings unless otherwise specified. The rationale behind this decision is discussed further in Supporting Information section III.

Vegetation structure surveys are carried out annually at the Glen

Finglas site by the James Hutton Institute, Scotland, United Kingdom. Three measurements of vegetation height (to the nearest 5 cm) are made at each of the 25 sample points (front, right and left to the observer), as well as 56 intermediate points, using a measurement stick marked with bands at 5 cm intervals (see Fig. 1 (right) for a visualisation of sample points). Vegetation density is also recorded at the same points (front, right and left density), by looking down the length of the measuring stick into the vegetation and recording the lowest 5 cm band that is visible (Douglas et al., 2008). To aid in analysis and to counter the potential to have imprecisely located the sampling points, the vegetation community at each point is recorded using the British National Vegetation Classification (NVC) (Rodwell, 1991).

We used climate data taken from the E-OBS 0.1° ×0.1° daily gridded European climate dataset (Cornes et al., 2018). This provides 7 climate variables collected using daily readings, which are then processed to generate 26 total climate metrics. The grid size at Glen Finglas is approximately 11.1 km × 7 km. Block E&F lies on the corner between four grid cells and the other two blocks also lie near the edge of their respective cells. In order to account for the coarseness of the data we have taken an average of the climate data over all of Glen Finglas and used it to study any landscape-wide effects, rather than trying to use coarse data to study the difference between the three blocks.

### 2.2 Analyses

#### 2.2.1 Cycle Analysis

When analysing the VSI time series, mean cycle amplitude is calculated as the mean of peak values minus trough values. Mean period is the mean of time between troughs and time between peaks. Max amplitude is defined as the largest difference between consecutive peak/trough values. Peaks are defined as local maxima within a neighbourhood of ±2 years. Troughs are likewise defined as local minima within a neighbourhood of ±2 years. Pearson’s correlation coefficients are used to test covariance, these were calculated using base R.

#### 2.2.2 Generalised Linear Mixed Model

To test for the effects of livestock grazing treatment on VSI and vole cycle amplitude, we used a generalised linear mixed model (GLMM) implemented in R using the GLMM Template Model Builder (*glmmTMB*) package (Brooks et al., 2017). Years where one or more sampling windows were missed (2005, 2017, 2020, 2022 and 2023) were excluded from the model to allow full factorial comparison. VSI was modelled as a binary outcome (presence/absence) using a binomial distribution with a logit link function. Due to the hierarchical nature of the experimental design, with multiple sample points within treatment plots, within larger experimental blocks, **Plot:SamplePoint** was included as a random intercept term, allowing us to capture both the within- and between-plot variation. Random slopes were initially tested; however, they were removed due to variance estimates being close to zero and causing convergence issues, suggesting that the additional complexity was unnecessary.

Fixed effects and their interactions were added sequentially following the original approach of Evans et al. (2006). Starting with a null model, we then added **Year, Treatment, Block** and **Season**, followed by higher-order interactions. The final model was selected using a combination of pairwise likelihood ratio tests (LRTs) and Akaike information criterion (AIC) comparisons. Interactions which did not improve model fit were excluded from the final model, including **Treatment:Season, Block:Treatment** and all third-order interactions and above. For full table of AIC and LRT results see Supporting Information section I. The final model selected was:

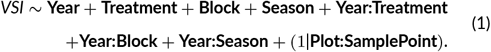

Although it was shown that the main effect of **Season** did not improve model fit based on LRT and AIC it was retained because the interaction of **Year:Season** strongly improved the model (it decreased the AIC more than any other effect), and our hierarchical model structure requires the inclusion of constituent main effects.

##### Model Diagnostics

Model assumptions were tested on the full model using the DHARMa package (Hartig, 2025), which simulates residuals in order to test uniformity (via QQ plots), dispersion, zero-inflation, outliers and temporal autocorrelation. Residuals were shown to be approximately uniform, with no significant deviations, and no evidence for over/underdispersion, zero inflation and influential outliers was found (see Supporting Information section I for plots and further details).

When checking temporal autocorrelation, residuals were first aggregated by year. The resulting ACF plots showed only weak correlations (none exceeding 0.4) and the Durbin-Watson test indicated no significant autocorrelation (DW = 1.4352, *p*= 0.23853). Random effect variances were checked and shown to be non-zero (≈ 0.5), confirming the hierarchical structure was appropriately accounted for.

Multicollinearity was evaluated using Variance Inflation Factors (VIFs) from the *car* package (Fox & Weisberg, 2019). Adjusted GVIF(1/(2^∗Df^)) values ranged from 1.33 to 5.51, with all but one term (**Year**) being below five. **Year** having an elevated VIF reflects its inclusion in all three interaction terms and is not considered problematic. Overall, diagnostic tests indicated that model assumptions were well satisfied and that the fitted GLMM was robust.

#### 2.2.3 Wavelet Analysis

Wavelet analysis is a powerful way of studying cyclic dynamics, in particular allowing us to observe dominant frequencies in the oscillations and how these vary temporally. A wavelet analysis was conducted following the approach of Torrence & Compo (1998) and Cazelles et al. (2008).

Specifically, we performed a continuous wavelet transform (CWT), decomposing our time series data into both time and period (scale). The CWT is defined as:

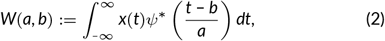

where *x*(*t*) is our original time series data, *ψ* is the Morlet mother wavelet, *a* is the scale (relating to both period and frequency) and *b* is the time shift. A Morlet wavelet is a complex sine wave with a Gaussian envelope, which can be stretched and shifted by time and period to match localised features of our time series data. The integral is used to calculate the “overlap” between the original data, *x*(*t*), and the scaled/shifted wavelet conjugate, *ψ*^∗^. The more that the wavelet matches the original signal’s shape for a value of *a* and *b*, the higher the value of the wavelet coefficient *W*(*a, b*).

The wavelet power spectrum (WPS) is defined as |*W*(*a, b*)|^2^, representing how much power is present at a given time and scale. This can be used to create heat maps, where the colour refers to where the periods of oscillation are particularly strong.

All wavelet analysis was performed in R using the biwavelet package (Gouhier et al., 2021).

#### 2.2.4 Mixed Effect Model

Linear mixed effect models were used to test the relationship between vegetation structure and VSI. The models were fitted using the *lme4* package in R (Bates et al., 2024). VSI was modelled as a function of both vegetation height and density, with the sample point included as a random intercept to account for spatial dependence. Both models assume that the residuals are normally distributed and that random intercepts capture between-point variability in VSI. The models are:

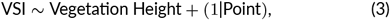

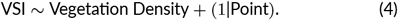

This analysis was performed twice. Once with the data collated across all grazing levels, and once split by individual treatment.

## 3 RESULTS

### 3.1 Relationship between grazing treatment and vole population cycle

There was a clear effect of treatment on VSI, apparent in Fig. 2, with the ungrazed plots (T4) having higher mean VSI than the grazed treatments (T1-T3). We observe that VSI across T2, T3 and T4 were more strongly correlated with each other than with T1, see Table 2. Temporal cycles in the VSI occur in all four treatments; however, increasing grazing pressure leads to smaller cycle amplitude. The cycle characteristics for all four treatments are shown in Table 3.

**TABLE 2.** Pearson’s correlation coefficients of VSI between livestock grazing treatments.

|  | T1 | T2 | T3 | T4 |
| --- | --- | --- | --- | --- |
| T1 | 1 | - | - | - |
| T2 | 0.754 | 1 | - | - |
| T3 | 0.766 | 0.822 | 1 | - |
| T4 | 0.711 | 0.873 | 0.861 | 1 |

**TABLE 3.** Characteristics of vole population wave.

| Treatment | Mean VSI | Mean Cycle Amplitude | Mean Period (Years) | Maximum Amplitude |
| --- | --- | --- | --- | --- |
| T1 | 0.0345 | 0.0506 | 3.35 | 0.133 |
| T2 | 0.0851 | 0.0974 | 3.28 | 0.225 |
| T3 | 0.0695 | 0.0716 | 3.23 | 0.214 |
| T4 | 0.1246 | 0.1183 | 3.38 | 0.412 |

**FIGURE 2.**
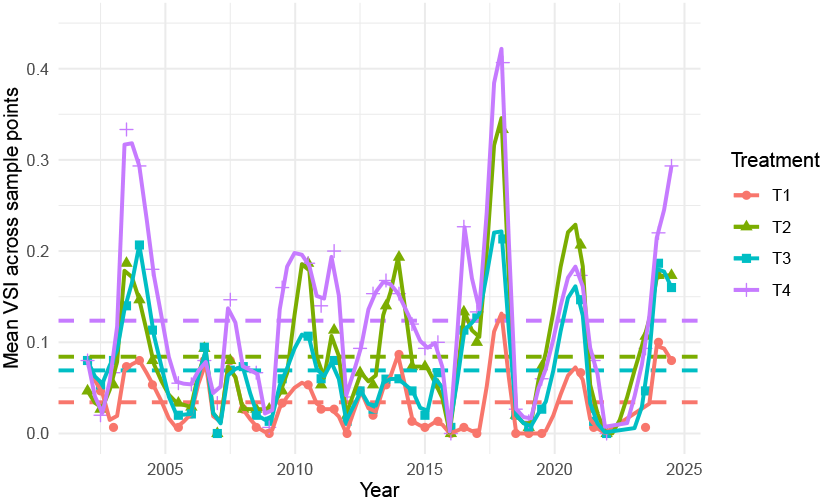
The time series of mean VSI (droppings) across sample points split by treatment (see Table 1). Average VSI is calculated as the mean VSI over all five quadrat throws for every sample point in each treatment. Solid lines show LOESS fits (span = 0.8) to aid visualisation. The dashed horizontal lines show the overall mean for each treatment over the entire course of the experiment, see Table 3 for treatment live-stock grazing densities.

A sequential generalised linear mixed model (GLMM) was performed to test the significance of multiple predictor variables on VSI, as in Evans et al. (2006). The final model, shown in Eqn. 1 and summarised in Table 4, retained all main effects and also included three two-way interaction terms. The inclusion of the **Year** factor in every one of these interactions shows the system’s strong temporal heterogeneity, suggesting that vole abundance is structured by long-term temporal dynamics, as to be expected in a cyclic system. In the Wald *χ*^2^ tests, the main effect of **Treatment** was not statistically significant; however, this does not imply that grazing treatment itself is not significant. Rather, the **Year:Treatment** interaction, which is significant, captures most of the biologically meaningful effects. In general the differences between treatment are consistent (T1 < T2 ≈ T3 < T4); however, the magnitude of these differences varies between which part of the cycle we are in (and hence varies by **Year**). Similarly **Season** as a main effect was eventually found to be significant in the final Wald *χ*^2^ test, but this effect is modest compared with the **Year:Season** interaction’s highly significance. This pattern is characteristic of a cyclic system, where there is no fixed seasonal difference within years, but rather a temporally varying relationship between seasons that shifts with the cycle phase. The inclusion of the **Year:Block** further shows significant spatial heterogeneity in vole dynamics over the experimental blocks. As a whole this GLMM provides strong evidence that grazing treatment has an important long-term impact on the amplitude and structure of vole cycle dynamics.

**TABLE 4.**
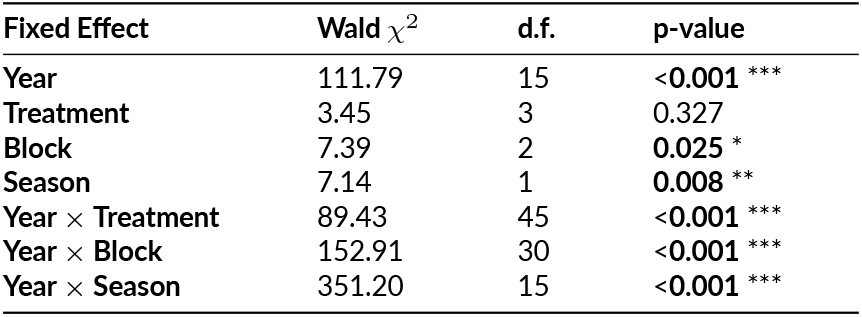
Fixed-effect terms from the final GLMM (Eqn. 1) testing VSI. The Wald *χ*^2^ statistic tests whether each predictor’s coefficients differ from zero conditional on all other terms. Degrees of freedom (d.f.) reflect the number of estimated parameters in each term, and the p-value indicates statistical significance, with p < 0.05 highlighted in bold.

### 3.2 Effect of experimental block

There was a clear effect of block (the three large, spatially separate, experimental areas) on VSI, with block A&B having significantly higher mean VSI than the other two areas, and block C&D having the lowest mean, as seen in Fig. 3. Again, temporal VSI cycles occur at approximately the same frequency in all three blocks, with likewise similar amplitudes.

**FIGURE 3.**
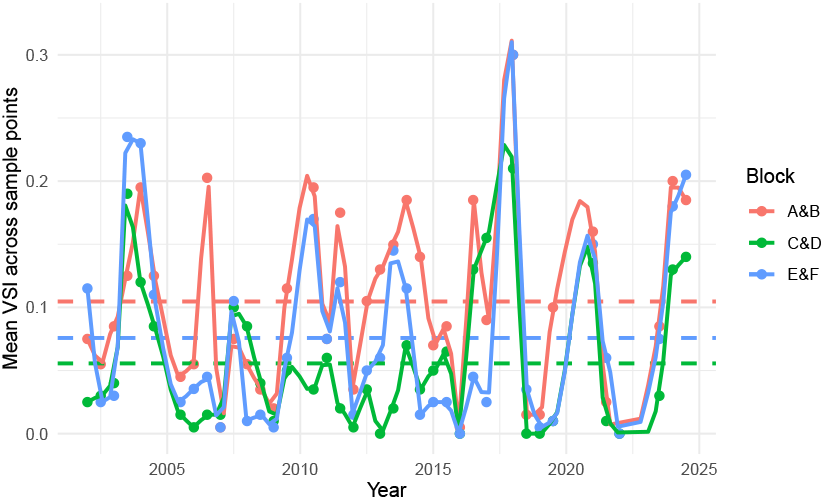
VSI by droppings over time grouped by block. Dashed lines represent the average VSI. Solid lines show LOESS fits (span = 0.8) to aid visualisation. Average VSI is calculated as the mean VSI over all quadrat throws for every points in each block.

### 3.3 Effect of vegetation on VSI

Two linear mixed effects models were performed to test the effect of vegetation structure on VSI. Both models returned significant results with *t*-values of 2.41 and 5.33 respectively, showing a positive relationship between vegetation height/density and VSI. The estimated effects were small in absolute terms (*β*_height_ = 0.00035 and *β*_density_ = 0.00155), but considering the mean VSI ranges from 0–1 while height ranges from 0–127 cm and density from 0–88 cm, this corresponds to biologically meaningful shifts across the observed gradients.

We then performed the same model analysis, but with the data split by treatment. Across all treatments, VSI showed little response to vegetation height, with only small, insignificant effects (|*t*| < 1.9). However, vegetation density was more strongly associated with VSI, with significant positive trends in T1 (*t* = 3.18, *p* < 0.01) and T2 (*t* = 3.37, *p* < 0.01). T3 and T4 exhibited much weaker, non-significant effects (*t* = 1.74 and *t* = 1.39). Random intercept variances were relatively small, indicating that VSI is not strongly tied to specific areas within plots, and rather varies over the years. See Supporting Information section II for full results.

To examine how vegetation structure has changed over the course of the experiment’s history, we show time series for both vegetation density split by treatment and block in Fig. 4 (a similar plot is available for vegetation height in Supporting Information section II).

**FIGURE 4.**
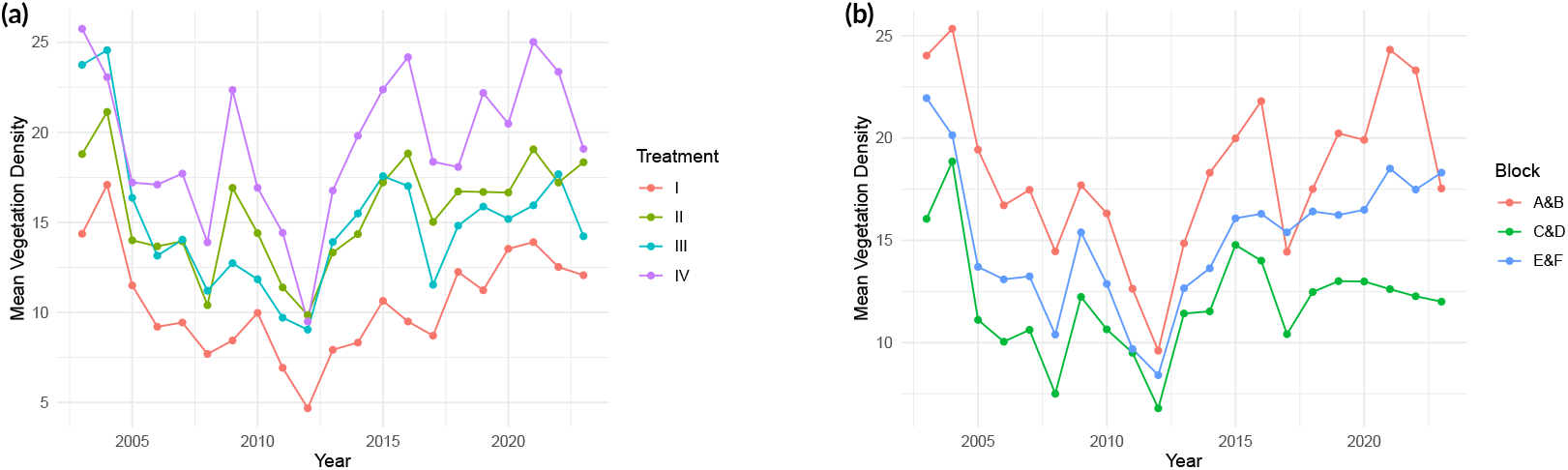
**(a)** Vegetation density time series by treatment **(b)** vegetation density time series by block. Average vegetation density is calculated as the mean vegetation density per point across each respective block and treatment

### 3.4 Collapse of vole population cycles

Wavelet spectral analysis was performed to examine the vole population cycles, firstly on the data as split into treatment. All four treatments give similar results and there is a band of high power corresponding to a period of approximately 3.5 years. The plots from this can be found in Supporting Information section II. The same wavelet spectral analysis was then performed for the data split into blocks instead of treatments, shown in Fig. 5. We again see the band of high power at around 3.5 years, showing that there is a significant 3.5 year period in the vole cycles. The same significant region at 1.4 years between 2016 and 2019 is still present, as well as all three blocks having some degree of noise at lower periods. However, for block C&D (bottom) we see a clear gap in the 3.5 year period between 2009-2016, indicating that there was a breakdown in the 3.5 year vole cycle in spatial block C&D. This cycle collapse is not seen in the other two spatial blocks.

**FIGURE 5.**
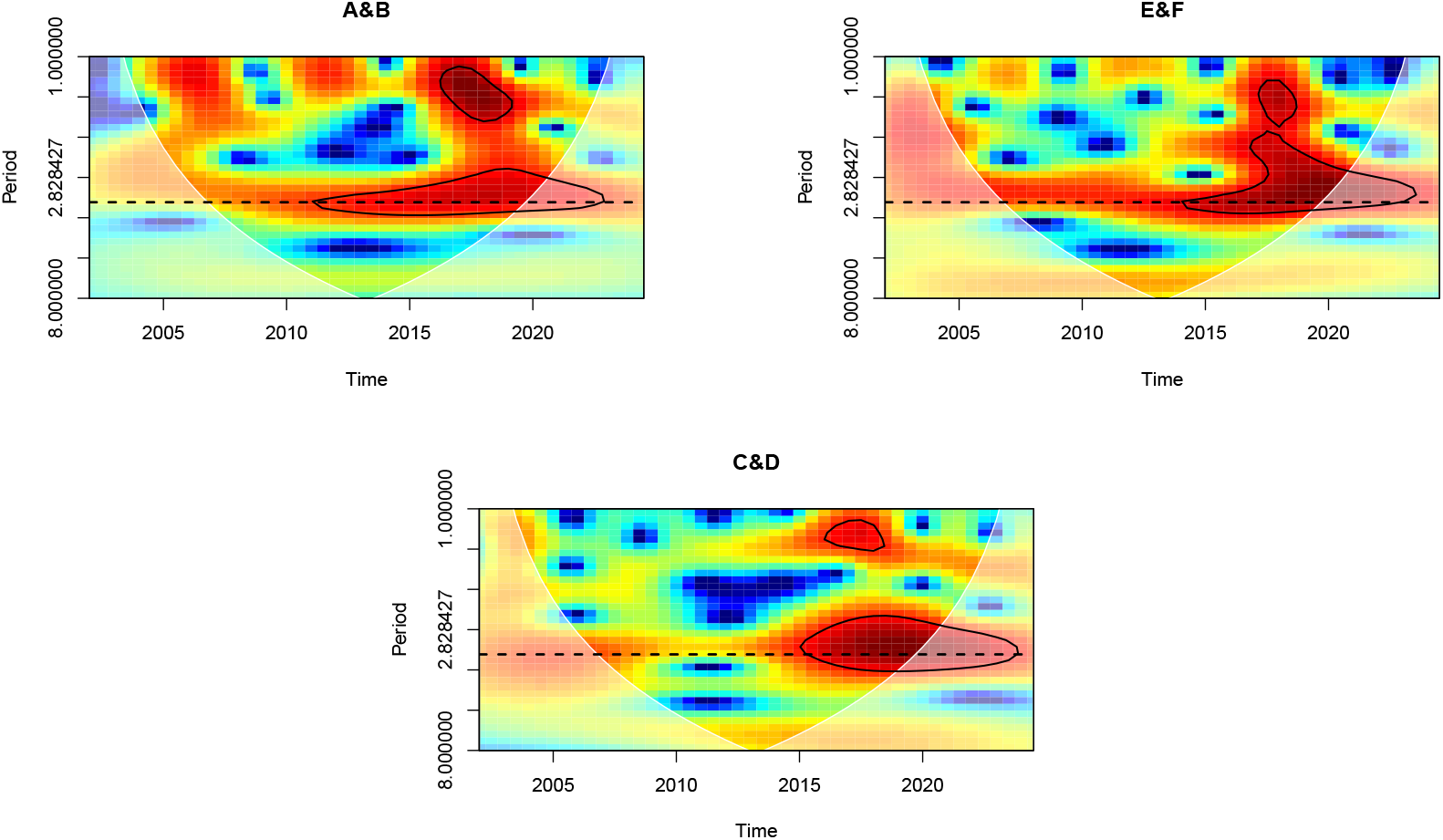
Wavelet Power Spectrum of each block. Y-axis is plotted on a log_2_ scale. The darker red shows a higher power period of oscillation, whilst the darker blue shows weak power oscillations. The white cone shows uncertainty due to the boundary effect. Black circles show a 95% significance level. Dashed line is at a period of 3.5 years.

We then checked to see if the lower mean VSI observed in block C&D is as a result of this collapse. We removed the 2009-16 period from the data and calculated the mean VSI again. With the dampened time-period removed, we see that block A&B actually sees a reduction in mean VSI (0.100 down from 0.104), whilst block C&D and block E&F both increase by about 0.01 (0.055 to 0.066 and 0.075 to 0.085 respectively). This shows that whilst the period between 2009-16 does account for some of the difference in mean VSI between blocks, it is not entirely responsible for this difference.

Studying the climate data in the years 2009-2016 we notice a trend across multiple metrics. Two of these metrics are highlighted in Fig. 6: average winter temperature and number of frost days (for more metrics see supplementary information). From Fig. 6 (left) we see that in 2010, near the beginning of this period of cycle collapse, there was an exceptionally cold winter. We see further, from Fig. 6 (right), that there were multiple years, including 2010, which had a larger number of frost days than in recent history. Other variables such as: winter rainfall, autumn temperature, mean yearly temperature, winter degree days minimum and maximum temperature and the yearly mean of minimum temperatures all showed similar patterns. However, note that some indicators such as wind-speed, rainfall, solar radiation and yearly mean of maximum temperatures showed no patterns corresponding to the cycle collapse. Further formal analysis of the climate data is presented in Supporting Information section IV.

**FIGURE 6.**
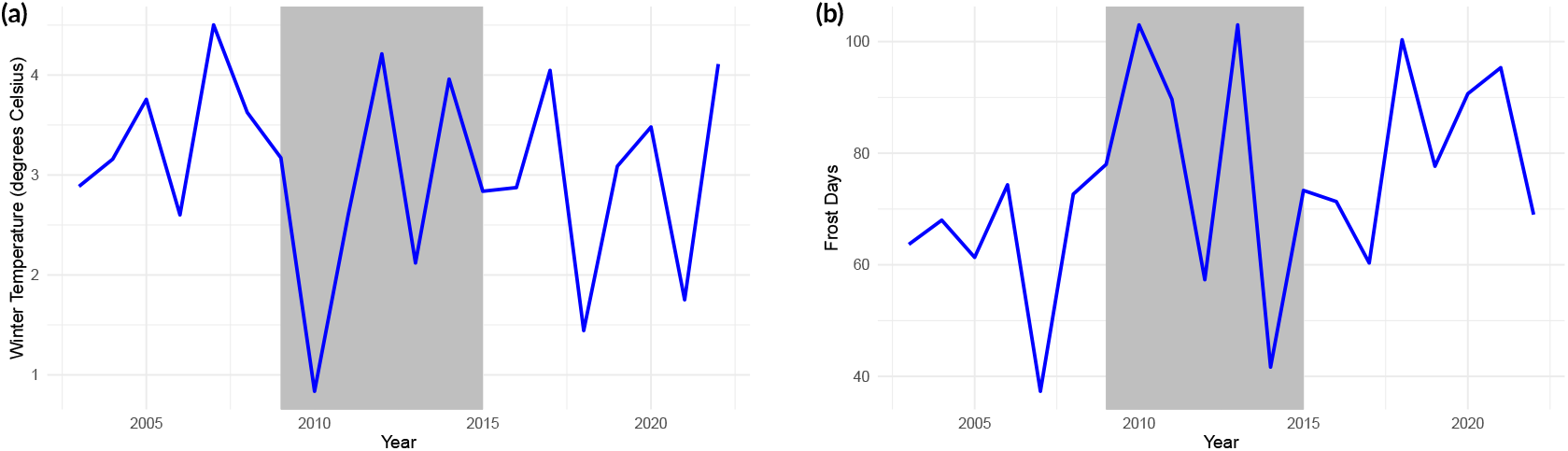
Time series of **(a)** winter temperatures and **(b)** frost days at Glen Finglas. The area between 2009-2015 is highlighted in grey to show where cycle collapse has been observed.

## 4 DISCUSSION

We have shown more heavily grazed plots have dampened oscillations with lower means whilst the ungrazed plots have significantly greater cycle amplitude and higher mean VSI. Oscillation period remained consistent across all four grazing treatments, with ~3.5 year cycles being dominant across the majority of the experiment. This suggests that the interaction between voles and large grazers is not responsible for driving vole cycles, but does influence the size of them. There was no significant variation between experimental blocks, except during the years between 2009-2016. In this period we have shown the presence of a localised cycle collapse in block C&D. Leading up to this cycle collapse we saw that 2009 and 2010 had unusually cold and long winters with above average snowfall. We also saw that vegetation density became especially low in the middle of this collapse. This suggests that perturbations, from both biotic and abiotic drivers, to vole populations, could lead to the destabilisation of vole cycles.

### 4.1 Relationship between grazing treatment and vole population cycles

We found significant effects of grazing treatment on vole population cycle dynamics, with highest cycle amplitude in ungrazed plots and lowest in intensively grazed plots. Whilst it is clear that treatments with less grazing result in higher vole populations and vice versa, the mechanism behind this relationship is less clear. There are multiple competing hypotheses which could likely explain this inverse correlation between grazing treatment and VSI. Firstly, competition with larger herbivores (i.e. direct effect) (Hansen et al., 1999; Steen et al., 2005). It is possible that small (e.g. voles) and large herbivores compete for food resources, therefore more livestock-heavy fields means more competition and hence fewer voles. It is therefore important to consider the grazing types of these large ungulate grazers and the foraging habits of voles. Voles mainly target fresh, green, shoots of softer recumbent grasses and flowers (Stenseth et al., 1977; Ferns, 1976; Evans, 1973; Hansson, 1971), similarly to sheep targeting newer plant growths and softer, more palatable plants (Cowlishaw & Alder, 1960; Rutter, 2006; Tóth et al., 2018). Cattle, however, are relatively indiscriminate in their grazing and tear large areas of turf up at a time (Cowlishaw & Alder, 1960; Rutter, 2006; Tóth et al., 2018; Hongo & Akimoto, 2003). This suggests that voles compete more directly with sheep than with cattle. However, our data shows slightly increased vole populations in areas grazed exclusively by sheep (T2) than in those subjected to mixed grazing (T3). The second hypothesis proposes that large herbivores indirectly influence vole habitat by altering its structure or composition. The ungrazed plots of T4 provide a much higher level of thicker vegetation cover than the heavily grazed T1 plots. The thick growth of vegetation in T4 means more habitat for voles to create their runs, dens and also provide cover from avian predators (Preston, 1990). The effect of vegetation height and density on terrestrial predators, such as specialised weasels and generalist foxes, is less clear however. It is likely that red foxes require a ‘goldilocks’ amount of vegetation for successful hunting in order to provide sufficient cover for ambush whilst not being blocked from reaching prey due to thick undergrowth (Laux et al., 2022), and for weasels there have been few studies but it is likely the dense vegetation is less of a barrier as they have evolved to travel through vole runs (Jędrzejewski et al., 1992). In a previous paper on this study, it was shown that fox activity was higher in the T4 plots; however, this was linked to these having higher vole populations (Villar et al., 2013). It is unclear whether this higher fox abundance led to a higher vole mortality rate due to predation. Whilst T1 (conventional stocking rate) leaves far less vegetation for breeding and shelter, it is important to note that there is still plenty of long, thick grass for shelter, just perhaps it is less effective or allows more predators access (Fig. 4 (left) and Supporting Information section II). Furthermore, differences in grazing style between cattle and sheep also means that T2 and T3 contain a different microhabitat structure. In general, cattle grazing leaves a more evenly grazed sward with fewer dense pockets of vegetation, whilst sheep leave undesirable areas alone, creating a mosaic structure. This has shown in particular to be the case for *Molinia caerula* dominated habitats (like Glen Finglas) as sheep selectively avoid *Molina* and preferentially graze other plants (Smith et al., 2014). However, cattle do create a larger variance in vegetation height, likely due to trampling effects (Evans et al., 2015). In Fig. 2 we see that T2 (sheep only) has a higher mean and peak VSI than T3 (mixture of cattle and sheep). As both of these treatments have approximately the same vegetation off-take, it is possible that voles benefit from the sheep’s grazing style more than they are hampered by the direct competition with sheep.

The final hypothesis which is first proposed in Massey et al. (2008), states that the interaction between large and small herbivores is mediated through plant silica defences. They provided clear experimental evidence that silica is used as a plant defence mechanism against herbivory (Massey et al., 2006), and furthermore that herbivory subsequently increases silica content in leaves (Massey, Roland Ennos, & Hartley, 2007). This is particularly important as we know that both grazing from large herbivores increases silica levels in vegetation (Massey et al., 2009), and that silica is disproportionately detrimental to voles compared to larger herbivores (Massey, Ennos, & Hartley, 2007). Massey & Hartley (2006) found that increased silica levels in grasses lead to: an increase in leaf abrasiveness, a smaller proportion of leaf consumed per vole, less nitrogen absorbed from grasses and ultimately a slower growth rate of voles when they are limited to a higher silica content diet. If this hypothesis were true we would expect that T1 would have the highest baseline silica content of all the treatments and T4 would have the lowest, as it has no extra grazing pressure from large ungulates (although deer can and do pass through the plots). Already field-work has shown that a key species increased under commercial grazing in T1 is *Nardus stricta*, which has a naturally higher silica content (Pakeman et al., 2019). It has been shown subsequently that it is mathematically plausible for this interaction between small herbivores and plant silica to drive vole population cycles (Reynolds et al., 2012, 2013); however, there has been no study of the system involving a large grazer also. This leaves a gap for future work in both testing whether there is a shift in baseline silica levels between treatments, and whether a mathematical model can be constructed to explain the interaction between treatment and vole cycle using silica.

In a recent paper by Lambin et al. (2025), it is suggested that delayed density-dependent recruitment is the driving force behind the vole population cycles, as opposed to survival (i.e. predation). Of these three aforementioned hypotheses, plant-silica defence acting as an interaction mediator would most fluently fit with recruitment driving cycles. Increased silica has been shown to slow down growth rates in young voles, but is unlikely to affect mortality significantly (Massey & Hartley, 2006), suggesting that as silica levels increase, vole recruitment would slow. A full study into grazing pressure during different cycle phases in a delayed density-dependent vole population model would provide further insight into this new hypothesis.

Despite all of these variations between treatments, the frequency of the VSI (and therefore population) oscillations remains remarkably consistent between treatments with the period always remaining around 3.5 years. From this we could hypothesise that the interaction with grazers does not produce cycles, but only influences the cycle amplitude and vole population size. However, it is worth noting that each experimental plot borders two to five other experimental plots as well as unmonitored land bordering the experimental plots, and whilst fences are in place to stop the movement of the large grazers, these fences provide no hindrance to the movement of voles. For the spikes in VSI (and therefore vole population) in 2003, 2017, 2020, and 2024, (Fig. 2) the T1 VSI exhibited a delayed response, remaining stable initially before increasing one time step later than the other treatments. It is possible then that in the local minima of the cycle, voles almost entirely disappear from T1 and only reappear when recolonising from adjacent plots during peak years. This may suggest that sufficient ungrazed vegetation may be a necessary condition for vole populations to cycle. There is some support for this as shown in Table 2 as it can be seen that T1 has considerably lower correlation than the other three treatments have with each other, suggesting that T1 is a unique case. Furthermore, in Fig. 2, the period between 2009 and 2016 is characterized by a noticeable dampening of the oscillations across all treatments. Here the oscillations we see so clearly in the rest of the plot become less sharp. This phenomenon has previously been observed in other vole population studies (Ims et al., 2008; Hörnfeldt, 2004; Hongo & Akimoto, 2003); however, like many of these other systems the cycle seemed to return to normal after just a few years (Brommer et al., 2010). We discuss this in more detail in 4.4.

### 4.2 Effect of experimental block

Although the ~5 km distance between experimental blocks is geographically minor, topographic variation within Glen Finglas (evident in Fig. 1 (left)) does introduce localised environmental differences. The hilly terrain results in differential exposure to sunlight and wind, particularly affecting block C&D. To account for these potential localised effects, we tested for a block effect to determine whether spatial stratification of the data was necessary.

When comparing the time series of VSI between blocks, Fig. 3, there is a hypothesis that the variation in mean VSI is to do with the change in environmental factors. If this were the case we would expect block E&F to be close to A&B in the vole activity levels; however, block A&B has significantly higher mean VSI (vole abundance). There is a possibility that this variation is a result of a factor not yet explored, such as vegetation composition or something not measured e.g. soil quality or plant nutrient composition. Differences between plots have not been studied since the original project report in 2005. This report showed A&B were the wettest of the plots and C&D are driest with E&F in between (Pakeman, 2025).

Perhaps the most striking difference between the blocks that we see in Fig. 3 is the period between 2009-16. Whilst the red and blue lines, for blocks A&B and C&D respectively, do not have the same clear wave formation as seen in the rest of the time series, there are still clear peaks in autumn 2010 and spring 2014 and a trough in spring 2012. However, in block C&D there is no discernible multi-year wave pattern at all between these years – indicating a localised, short-term cycle collapse. We discuss this in greater depth in section 4.4.

There is strong synchronisation of vole cycle frequency between blocks, except for the collapse of cycles between 2009-16 in block C&D. We removed the period of cycle collapse from the data to examine whether this accounted for block C&D having the lowest average VSI. Whilst this does bring the average VSI in the three treatments slightly closer together, it fails to account for much of the variation between the three blocks. This shows that there is a significant effect of block on the measured VSI. However, it remains unclear as to whether this variation is due to climate effects, vegetation differences or something else entirely.

### 4.3 Effect of vegetation structure on VSI

The relationship between vegetation structure and VSI is strongly related to the relationship between treatment and VSI, as there is naturally a significant relationship between vegetation and grazing intensity.

Fitting of mixed effect models for vegetation height and vegetation density, it can be seen that when the data was viewed with all treatments together, both height and density were a significant variable in VSI. However, when split into treatments, only density in T1 (conventional stocking) and T2 (low level sheep only) were significant. These results suggest that whilst overall voles prefer more high and dense vegetation, it is only large changes to grazer stocking that make a difference - small changes in microhabitat across a treatment field were inconsequential. In treatments T3 and T4, it is likely that the vegetation is sufficiently dense everywhere to provide enough cover and food for voles, hence only in T1 and T2 (where vegetation is less dense overall) does density become significant.

The mean vegetation height has been consistently increasing across all experimental blocks and treatments, with treatments with less grazing pressure having higher vegetation (Pakeman et al., 2019). The overall trend is quite different for vegetation density (Fig. 4). The time series of vegetation density does not grow in a linear fashion like vegetation height, instead we see initial drops in vegetation density across all blocks (and treatments) before recovering up towards its pre-experimental levels. It is currently unclear as to whether this is due to the onset of experimental treatments, or if it is a landscape-wide trend. Although perhaps the most likely explanation is the two cold winters just prior to this, as shown in Fig. 6

We know that it takes a long time to observe community-level changes in vegetation (Pakeman et al., 2019). As voles are the main cause of seedling and sapling destruction, is the elevated presence in the T4 (ungrazed) plots contributing to this slowed ecological succession? Furthermore it is known that as the habitat changes from a *Juncus* and *Molinia* dominated acid grassland into a young forest, it becomes less suitable for field voles. Perhaps the combination of these two effects will create a bifurcation point where, once enough of the grassland becomes forest, it changes from slow change to a sudden shift of habitat due to the suppression of voles destroying young trees and seeds. Only one plot in the whole experiment - A4 - has reverted from grassland to young woodland, and this change did appear to happen rapidly. Further studies on vole destruction of saplings in ecological habitat succession would be needed to provide further insight into this observation.

### 4.4 Collapse of vole population cycles

Due to there being a number of recorded cycle collapses (Cornulier et al., 2013; Hörnfeldt, 2004; Hörnfeldt et al., 2005; Ecke et al., 2017), and subsequent recoveries over recent years, we checked whether a similar collapse occurred in our dataset and, if so, what factors may have contributed. As discussed above, we do observe a potential disruption in cyclic dynamics between 2009 and 2016, as shown in Fig. 2 and Fig. 3. The wavelet power spectrum (WPS), (see Supporting Information section II), suggests that for each treatment over the last 20 years there has consistently been the expected ~3.5 year *Arvicolinae* cycle (Krebs, 2013). The wavelet power spectra for each treatment show no evidence of population cycle collapse.

If we consider the WPS by experimental block, Fig. 5, we observe a break from the previous pattern: blocks A&B and E&F consistently have the 3.5 year cycle; however, for block C&D there is a significant gap in this between 2009-2015 - where the observed collapse was. So instead of a landscape wide cycle breakdown event, we instead observed a highly localised, small scale collapse - contrary to what other previous authors have noticed (Ims et al., 2008; Hörnfeldt et al., 2005). Further experiments will be necessary to elucidate the underlying mechanisms, but these findings raise several important questions for future research. One such question is whether subtle environmental differences between blocks are sufficient to disrupt vole population cycles temporarily.

Potential climate impacts are observable in our data. Whilst not consistent over the entirety of the 2009-2016 period - it is still clear to see from Fig. 6, that at the beginning of the cycle breakdown (2009 & 2010) there was a period of unusually low temperatures. Winter in 2009 and 2010 was characterised by deep snow which took a long time to melt. Snow and frost at the end of 2010 also came quite early. Some other climate metrics (such as precipitation, wind-speed and solar radiation) did not show any noticeable changes; however, these metrics do not directly relate to winter temperatures. As vole numbers are already lower going into the winter months when they reduce their activity, a particular cold winter could have a larger effect on the overall population as there will be fewer voles left to reproduce when spring arrives.

It is very likely that the interaction between large herbivore grazers and voles, and potentially the mechanism behind the cycle collapse, is an indirect interaction, mediated through vegetation. We see that whilst vegetation height remains relatively consistent in the collapse period (2009-16) - with vegetation density seeming to decline post 2009 (Fig. 4), possibly due to the cold winters of 2009/10 and 2010/11 (Fig. 6). Block C&D has the lowest vegetation density and height, partially due to C&D having the lowest dominance by *Molinia* and instead containing more *Nardus* patches, making it the most exposed to the cold and wind, whilst also likely containing less quality food (Pakeman et al., 2019). During cold winters, snow on well established *Molinia* tussocks could potentially insulate voles from freezing temperatures (like those seen in 2009-11), as seen in Fennoscandian voles (Haapakoski & Ylönen, 2013). *Nardus*, however, is unlikely to perform this same role as it does not form thick tussocks like *Molinia*. Unfortunately the E-OBS climate data does not measure snow cover so it is hard to say how much of an effect this has.

It is also true that block C&D are situated on the highest and steepest slope of all three plots, meaning they potentially receive an even larger effect from wind and weather. Pairing this with the low vegetation density and height of poorer quality cover leads us to hypothesise that block C&D lacks sufficient habitat and enough quality food in order to shelter voles from an abiotic driver (such as poor winter weather) causing a small perturbation in vole abundance. Figure 3 also shows that just before the cycle collapse in C&D, the VSI numbers were very low as the cycle was right in the bottom of a trough. The period of time in which we observe this cycle collapse was around early 2009 until late 2016 - almost exactly two full periods of the wave. A small perturbation in already low population numbers could potentially be what lead us to a collapse of the cycle for two full periods. Further experimentation comparing the blocks specific microclimate as well as voles’ reaction to environmental shock is needed in order to test this hypothesis.

There are other possible explanations for the change in the robustness of vole population cycles. Data shows the existence of a global 3.5 year temperature cycle which is now being affected by anthropogenic warming effects. If this temperature cycle is connected to the 3.5 year population cycle then there is a possibility that global changes in climate could impact voles directly. This discussion is explored further in Supporting Information section V.

## Conclusion

With the warming of global climate and acceleration of ecological degradation, it is important to understand how vole populations vary, as they can be detrimental to reforestation efforts - such as the plantation of a new broadleaf forest being carried out by the Woodland Trust at Glen Finglas. Our work shows there is a clear negative effect of increased live-stock grazing to vole abundance and cycle amplitude. Cycles were still consistently present and of a ~3.5 year period across all grazing treatments, showing that the interaction between voles and large-grazers is not the driver of vole cycles. We have also shown that between the years of 2009 and 2016 there was a substantial cycle collapse in the vole population oscillations; however, this breakdown was highly localised in only one of the three experimental blocks. It is likely that this collapse was caused by perturbations from biotic and abiotic drivers at the trough in the vole’s cycle. In order to further understand the drivers behind the vole cycles and the factors involved in their collapses, further research is needed into the mediators of vole-ungulate interactions. Perhaps the most promising avenue of further research is to test whether plant-silica defences can act as this interaction mediator, as multiple results from this paper could be explained by such a mechanism.

## Abbreviations

VSI: Vole Sign Indices;

## Acknowledgements

We thank The Woodland Trust, Scotland and Hamish Thomson for permission to use the Glen Finglas Estate. We also thank Debbie Fielding, David Riach, Aifionn Evans and other staff at JHI who have assisted with the vole and vegetation surveys and other data collection over the experiment.

## Notes

Funding Information This research was supported by the ONE Planet DTP and the Natural Environment Research Council (NERC) as part of MD’s PhD, supported by (Grant Number [NE/S007512/1]). LEW acknowledges support from NERC Knowledge Exchange Fellowship (Grant Number [NE/X000478/1]).

### Competing Interest Statement

The authors have declared no competing interest.

