## Supporting Information for "The impacts of grazing pressure by large herbivores on short-tailed field vole (*Microtus agrestis*) population cycles: results from a long-term upland experiment"

### I. GLMM TEST PLOTS AND RESULTS

The GLMM was first fitted using a sequential method as described in Section 2.2.2, the results from that process are presented in the table below. It was decided that **Block:Treatment**, **Treatment:Season** and **Year:Season:Treatment** would be excluded due to either their AIC, LRT or both.

TABLE I: Table showing change of Akaike information criterion (AIC) after each term is sequentially added as well as whether the likelihood ratio test (LRT) of each new term being added was significant (<0.001 ***, <0.01 **, <0.05 * , <0.1 ·, >0.1 n.s.). AIC in bold indicates low decrease/an increase and hence insignificance.

| Term Added | d.f. | AIC | LRT significance |
|---|---|---|---|
| Null Model | 2 | 6182 | n/a |
| Year | 17 | 5901 | *** |
| Treatment | 20 | 5834 | *** |
| Block | 22 | 5803 | *** |
| Season | 23 | **5804** | n.s. |
| Year:Treatment | 68 | 5792 | *** |
| Year:Block | 98 | 5667 | *** |
| Block:Treatment | 104 | **5666** | * |
| Year:Season | 119 | 5095 | *** |
| Treatment:Season | 122 | **5100** | n.s. |
| Year:Season:Treatment | 167 | **5122** | * |

Once the final model was fitted as described in Eqn. 1, DHARMa was used to test the model assumptions. The DHARMa tests for uniformity, dispersion, zero inflation, outliers and temporal autocorrelation were caried out first.

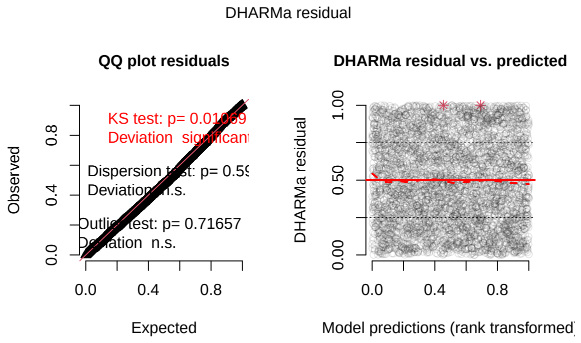


FIG. 1: DHARMa plot for uniformity (QQ plot).

Note that whilst the Kolmogorov-Smirnov (K-S) test in Fig. 1 shows significant deviation, this is common in large datasets, as with many observations even that smallest deviation can lead to the K-S test showing significance. As the other tests show no significance in deviation and the plot looks straight, it can safely be discounted.

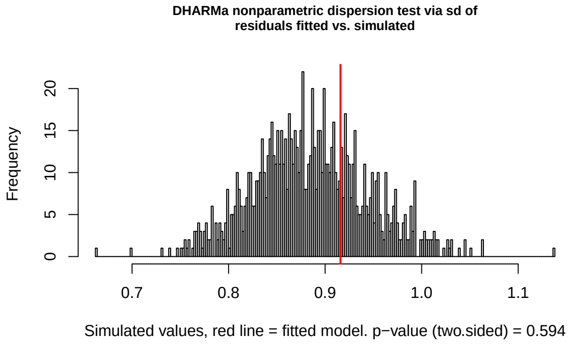


FIG. 2: DHARMa plot for dispersion.

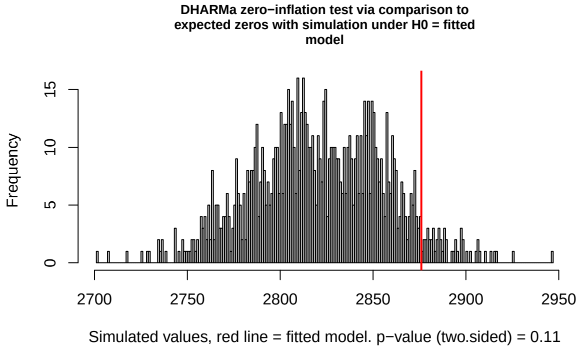


FIG. 3: DHARMa plot for zero inflation.

In both Fig. 2 and Fig. 3, the normal distribution shape with no skew shows there is no significant over/underdispersion or zero inflation. In Fig. 3 the predicted mean significantly above observed mean shows that the model predicts slightly more zeroes than the data shows; however, the effect is not significant. Similarly in Fig. 4 a relatively uniform distribution shows that there are no significant outliers.

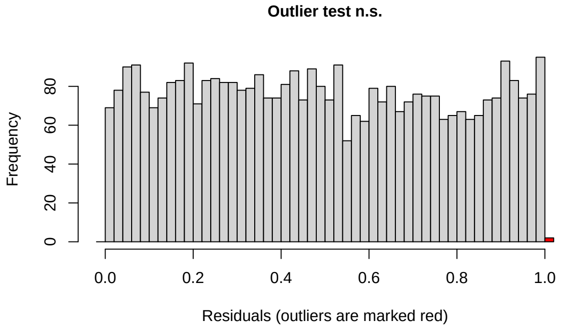


FIG. 4: DHARMa plot for outliers.

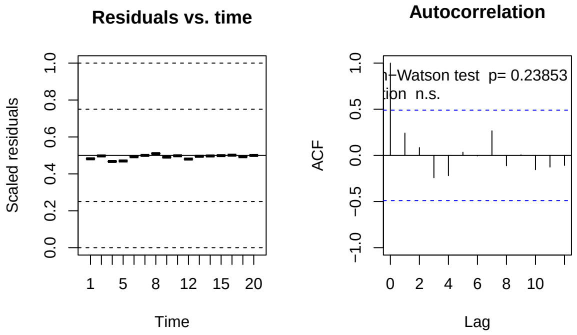


FIG. 5: DHARMa plot for temporal autocorrelation.

In Fig. 5, the residuals over time are approximately uniform at 0.5 which suggests no significant autocorrelation. The ACF plot (right) shows that for t>0 the level of correlation is well below the blue dotted significance bars.

We then tested whether any systematic structure remained unexplained by the model, we plotted the simulated residuals against key predictors again using DHARMa.

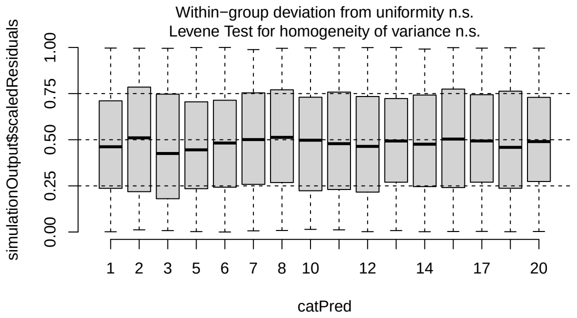


FIG. 6: DHARMa simulated scaled residuals plotted against Year

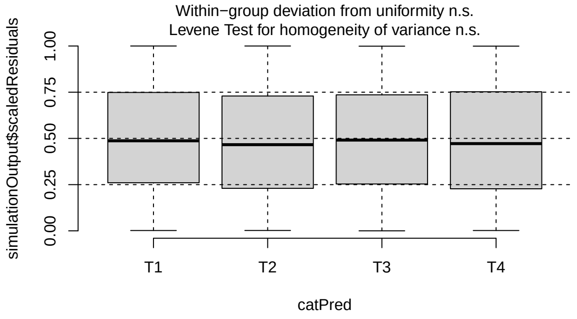


FIG. 7: DHARMa simulated scaled residuals plotted against Treatment

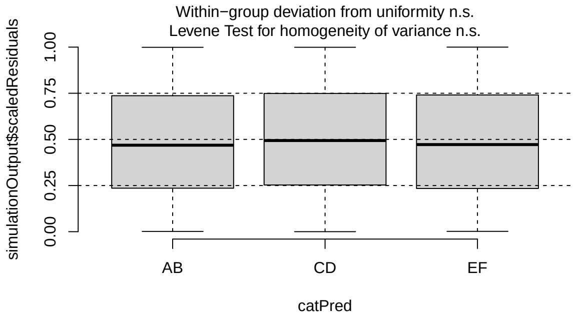


FIG. 8: DHARMa simulated scaled residuals plotted against Block

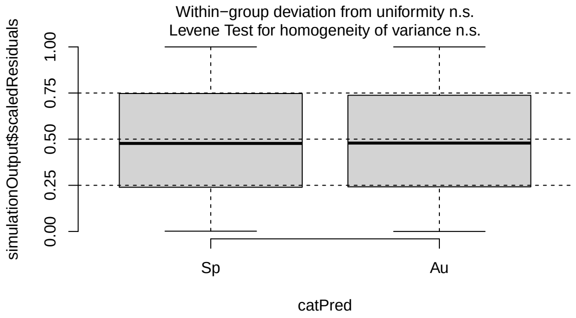


FIG. 9: DHARMa simulated scaled residuals plotted against Season

For all of these plots there appeared to be no apparent trends or deviations, indicating that the fixed and random effects included in the model adequately accounted for temporal and spatial variation.

Next random effects were tested. **Plot:SamplePoint** was included in the model as a random intercept, the estimated standard deviation of these random effects was 0.55, indicating moderate variation amongst plots and sample points even after controlling for the other fixed effects. The random intercepts ranged from -0.95 to 1.25 on the logit scale, suggesting that some location supported higher than average VSI, whilst others contained lower VSI; however, no extreme spatial outliers were detected. See Fig. 10 for a plot to help visualise this.

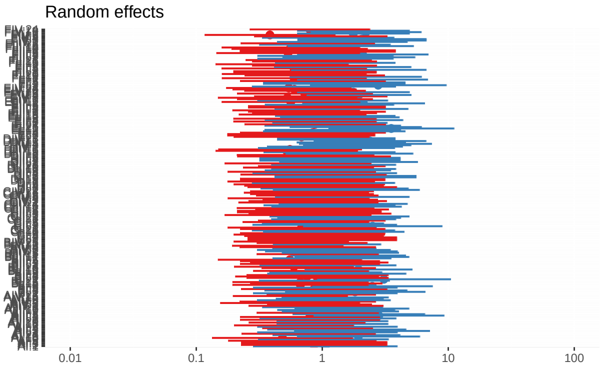


FIG. 10: Visualisation of the random intercepts from **Plot:SamplePoint**.

Finally mulitcollinearity was checked using VIF as discussed in Section 2.2.2. The results from the VIF are in the table below:

TABLE II: Table showing variance inflation factors (VIFs). The higher the VIF value the more colinearity there is between a given term and the other effects in the model. Generally a VIF $< 3$ is considered to be "excellent" and VIF<5 is "good".

| **Fixed Effect** | **GVIF** | **d.f.** | **GVIF**$^{1/(2*\mathbf{d.f.})}$ |
|---|---|---|---|
| Year | 1.7e22 | 15 | 5.51 |
| Treatment | 6.23e3 | 3 | 4.29 |
| Block | 4.4e2 | 2 | 4.58 |
| Season | 21.6 | 1 | 4.65 |
| Year:Treatment | 1.27e23 | 45 | 1.81 |
| Year:Block | 2.90e7 | 30 | 1.33 |
| Year:Season | 1.06e9 | 15 | 2.00 |

#### II. LINEAR MODEL RESULTS

On the following page is a table summarising all the results from the 10 linear models ran in R using the lme4 package [1]. Each model was checked for residual normality, homoscedasticity and outliers, as well as checking that the random effects were justified.

Long term analysis suggests that whilst there was an initial increase in VSI in some plots due to the inception of the treatment, this trend stopped within the first 4 years of the experiment. This is demonstrated in Fig. 11, where the inset plot shows the first four years of the experiment and the apparent initial increase of VSI across treatments which was observed in [2].

There is a noticeable change in the trend of rapid VSI increase in treatments T2, T3 and T4. For the first 3 years of the experiment it was believed that vole populations were growing substantially in these treatments; however, with the full 23 year dataset we can see that this trend does not hold (Fig. 11). There are again two possible reasons for the initial results showing a year-on-year VSI increase. Firstly that the implementation of the treatments lead to a rapid population growth in the ungrazed T4 plots, which spilled over

TABLE III: Table showing full results from the 10 linear mixed models ran. T is the treatment tested. RI SD is the random intercept standard deviation. Res SD is the residual standard deviation.

| **T** | **Predictor** | **Estimate** | **Std. Error** | ***t*-value** | **Significance** | **RI SD** | **Res SD** | **Interpretation** |
|---|---|---|---|---|---|---|---|---|
| All | Height | 0.00035 | 0.00015 | 2.41 | $p<0.05$ | 0.0459 | 0.1090 | Significant Positive Effect |
| All | Density | 0.00155 | 0.00029 | 5.33 | $p<0.001$ | 0.0426 | 0.1090 | Strong Positive Effect |
| T1 | Height | 0.00027 | 0.00017 | 1.52 | n.s. | 0.0202 | 0.0627 | Weak Positive - n.s. |
| T2 | Height | 0.00057 | 0.00030 | 1.91 | $p\approx 0.06$ | 0.0327 | 0.1166 | Marginal Positive Effect |
| T3 | Height | -0.00012 | 0.00029 | -0.43 | n.s. | 0.0360 | 0.1087 | No Relationship |
| T4 | Height | 0.00036 | 0.00035 | 1.02 | n.s. | 0.0412 | 0.1351 | No Relationship |
| T1 | Density | 0.00140 | 0.00044 | 3.18 | $p<0.01$ | 0.0178 | 0.0626 | Significant Positive Effect |
| T2 | Density | 0.00196 | 0.00058 | 3.37 | $p<0.01$ | 0.0327 | 0.1158 | Significant Positive Effect |
| T3 | Density | 0.00100 | 0.00057 | 1.74 | $p\approx0.08$ | 0.0329 | 0.1088 | Weak Positive Trend |
| T4 | Density | 0.00088 | 0.00063 | 1.39 | n.s. | 0.0401 | 0.1352 | No Relationship |

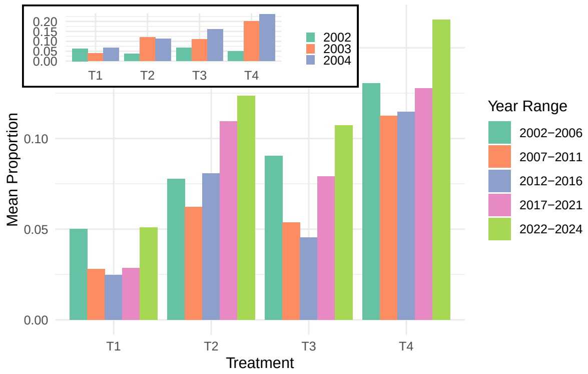


FIG. 11: A comparison with results from 2006 [2], contrasting early short-term changes to long-term trends. Inset plot shows treatment vs VSI for the first 3 years of the experiment. Main plot shows treatment vs VSI split into four year groups for the remainder of the experiment.

into neighbouring plots. Or perhaps instead it is simply that the experiment was started in the trough of a vole cycle, and the increase seen was a natural part of the oscillations. This is more clearly observed from the VSI by treatment time series, Fig. 2 in the main text.

Wavelet spectral analysis was performed to examine the vole population cycles, firstly on the data as split into treatment shown in Fig. 13. For all four treatments there is a band of high power corresponding to a period of approximately 3.5 years. There also appears to be a period of 1.4 years ($\pm$ 0.2 years) between the years 2016-2019. We see that T1 (top left) has considerably more noise at lower periods than the other three treatments. We also note that there is no discernible breakdown of the 3.5 year cycle between 2009-2016 for any treatment.

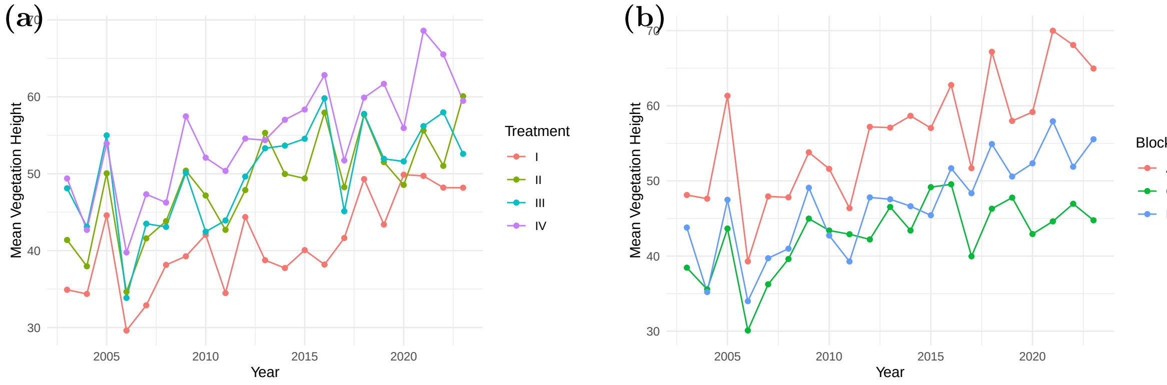


FIG. 12: Time series of mean vegetation height over Treatment and Block. Both are calculated as the mean of front, right and left vegetation height/density across all points in given treatment.

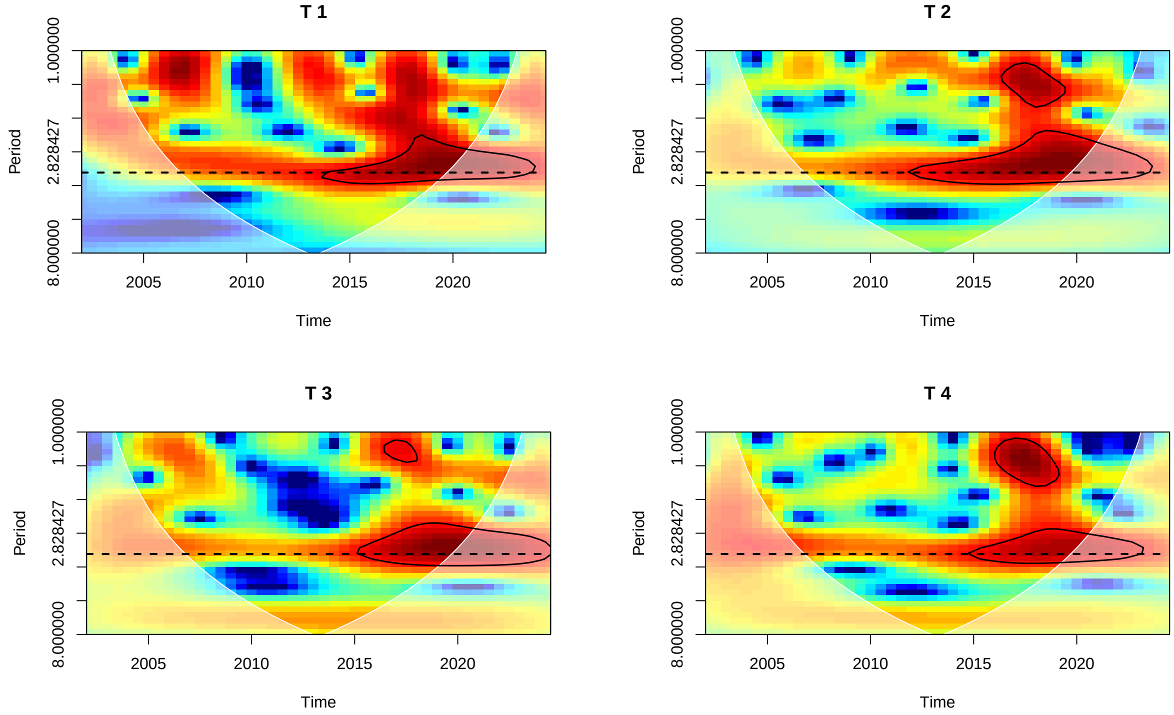


FIG. 13: Wavelet Power Spectrum for each treatment. Y-axis is plotted on a $\log_2$ scale. The darker red shows a higher power period of oscillation, whilst the darker blue shows weak power oscillations. The white cone shows uncertainty due to the boundary effect. Black circles show a 95% significance level. Dashed line is at a period of 3.5 years.

#### III. VOLE SIGN INDEX METHOD

When considering VSI as a proxy for vole population abundance, it is important that we understand how using each individual vole sign index affects our results. In order to get a better understanding of this we have plotted VSI over time, separated by vole sign index used, in Fig. 14.

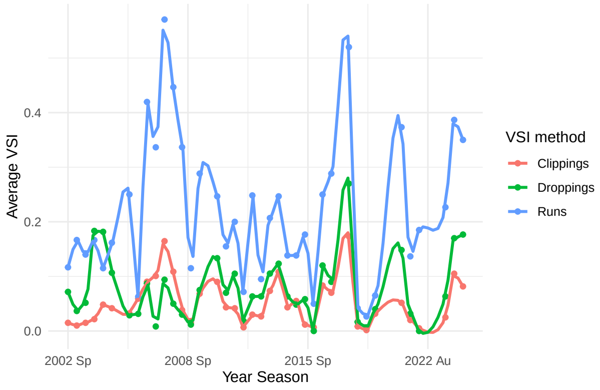


FIG. 14: A plot showing the comparison of VSI methods for all treatments. Solid lines show LOESS fits (span = 0.8) to aid visualisation. Average VSI is defined as the mean value of given VSI type found over all quadrat throws for each measured point over all treatments and blocks.

We see that runs are by far the most observed method of VSI, with both runs and droppings remaining approximately equivalent. We also see that all VSI methods used retain the same cycle patterns, and even smaller details are also consistently captured by all 3 VSI methods.

We also analysed how finding certain VSI affects how often we find other VSI - particularly in the context of runs as these were the most common VSI. We saw that if both clippings and droppings are present then we found a run 97.5% of the time. If there was either a dropping or clipping present then we still found a run 74.7% of the time. However, conversely if a run is found there is only a 35% chance of finding either a dropping or clipping and an even smaller 12% chance of finding both. If we believe that clippings and droppings are the more accurate VSI, then we can suggest that runs as a VSI have a good sensitivity but a bad specificity. This is also supported in the literature, where it is stated that without traces of the other two indicators it is hard to tell when the run was last used as they can stay around for a long time, so runs are inconsistent to be used as a main vole sign index [3, 4]. Initially we thought perhaps that the presence of large ungulate grazers caused the runs to be trampled - and hence disappear more quickly; however, tests showed that there was no significant interaction between treatment and runs, and we still saw the same sharp drops in the ungrazed patches. From all of this evidence we decided that runs were likely giving too many false positives so we decided not to use this VSI method in analysis.

Comparing clippings and droppings we see that if we find a clipping then there was a droppings present 51% of the time; however, if a dropping was found then only 35.1% of the time we found a clipping. When either clippings of droppings were found, only 26.2% of time was the other VSI also found. So despite both having similar looking time series, they are actually not found together often.

There was an option therefore to use a linear combination of both droppings and clippings in order to create a more reliable VSI. However, it was instead decided to use only droppings for three main reasons. Firstly the previous papers on the Glen Finglas experiment have largely focused on only droppings, so continuing this allows us to more directly compare results. Secondly using droppings as a singular VSI method means that it will likely be easier to compare our work to the wider literature as many other papers use only one vole sign index. Lastly the scientists who completed the data collection for this part of the experiment felt they had significantly more confidence in the reliability of the dropping data than the clippings data.

#### IV. IMPACT OF CLIMATE VARIABLES

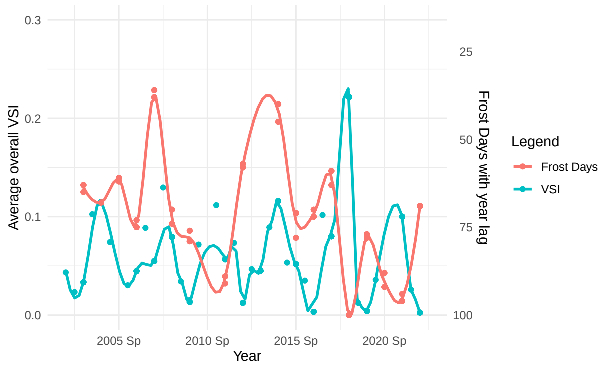


FIG. 15: VSI vs Frost days from previous winter. Average overall VSI is the mean value of VSI across all quadrat throws for every point for all treatments and blocks. Frost days are plotted on an inverted scale from 100 to 0.

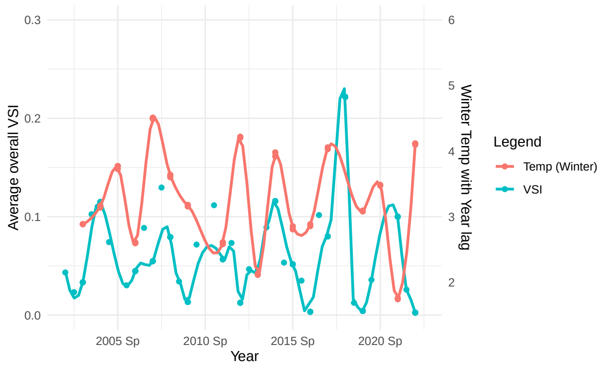


FIG. 16: VSI vs winter temp. Average overall VSI is the mean value of VSI across all quadrat throws for every point for all treatments and blocks.

We noticed that there appeared to be a negative correlation between the recorded frost days and VSI, with spikes in VSI in 2008, 2014, 2017 and 2021 seeming to match up well with particularly low frost days as seen in Fig. 15. However, Fig. 16 shows that this did not line up so well with mean winter temperature, which is strongly correlated with frost days. We then performed two linear regression tests, split by the spring and autumn measurements:

$$\text{VSI} \sim \text{Frost Days} + (1|\text{Treatment})$$
$$\text{VSI} \sim \text{Frost Days} + (1|\text{Block})$$

The results for these models are plotted in Fig. 17 and Fig. 18. As can be seen there was a slight negative correlation; however, neither model showed that this correlation was significant.

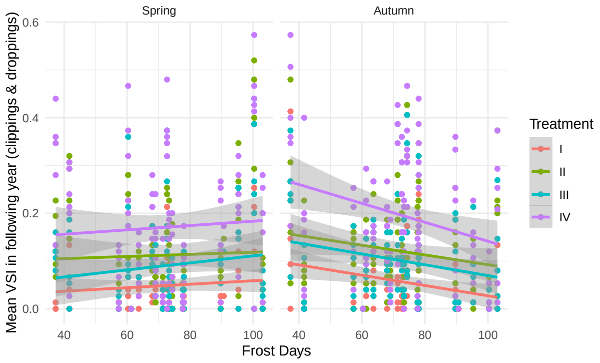


FIG. 17: Linear relationship between frost days and VSI by treatment.

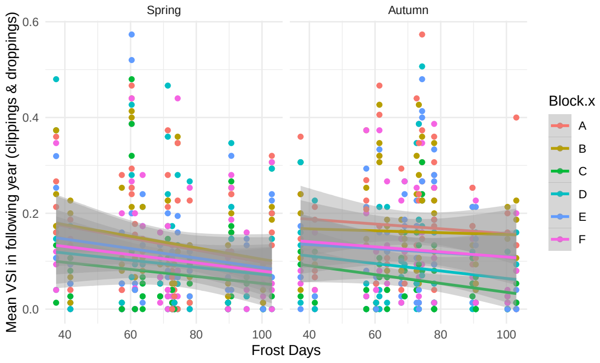


FIG. 18: Linear relationship between frost days and VSI by block.

We showed that, despite the visually promising results in Fig. 15, there was no significant correlation between frost days or winter temperature (or in fact any temperature metric) on VSI.

#### V. CLIMATE CYCLES AS AN INFLUENCE ON VOLE CYCLES

Solar cycles of 10-12 years are well known and documented [5]; however, the Earth actually has many other oscillating climate effects, such as El Niño-Southern Oscillation (2-7 years), Pacific Decadal Oscillation (20-30) years amongst others. Using data from NASA and performing a simple FFT we can see more of these (Fig. 20).

In Fig. 20 we can see a 3.5 year cycle occurs in global temperature over the last 150 years. This has been documented by climate scientists, but the forcing behind this isn't abundantly clear [6].

This data, however, is at a global scale and the 3.5 year temperature cycle is therefore not consistent at a local scale - take for example the Glen Finglas climate over the last 20 years in Fig. 21.

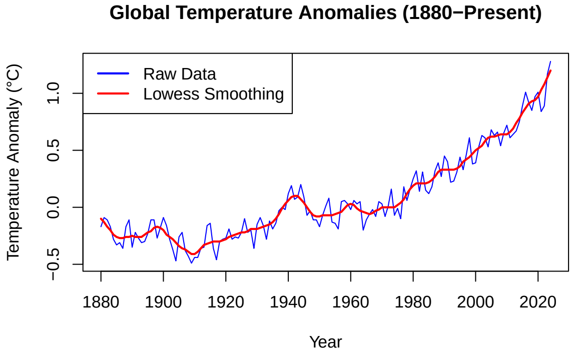


FIG. 19: Time series of global land-ocean temperature anomalies. Data source and credit: NASA's Goddard Institute for Space Studies (GISS).

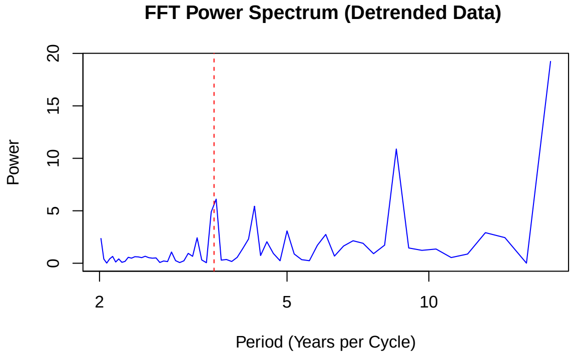


FIG. 20: Fast Fourier Transform (FFT) of global land-ocean temperature anomalies. Red-dotted line at 3.5 years. Data source and credit: NASA's Goddard Institute for Space Studies (GISS).

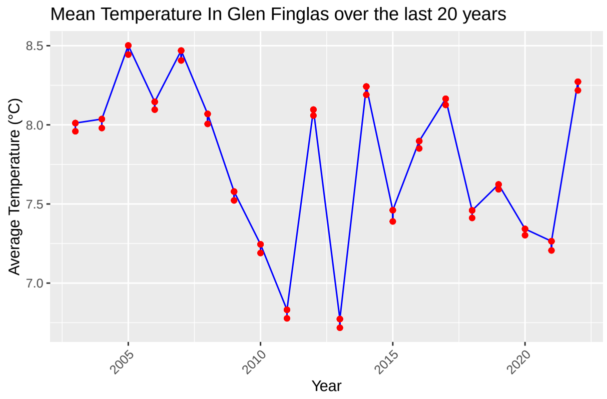


FIG. 21: Time series of Glen Finglas temperature.

Although the FFT in Fig. 22 picks up an oscillation of between 3.75-4.25 years, it is clear from the time series data in Fig. 21 that on this short scale there is no clear 3.5 year temperature oscillation to match with the VSI oscillations.

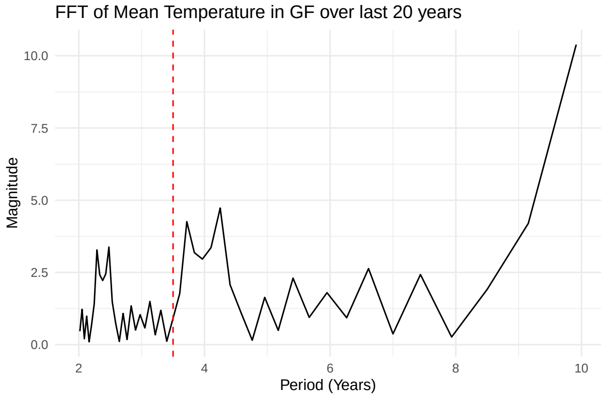


FIG. 22: Fast Fourier Transform (FFT) of detrended Glen Finglas temperature. Red-dotted line at 3.5 years.

Vole cycles are a global phenomena; however, outside of the northern hemisphere they don't necessarily share the same 3.5 year trend [7]. Furthermore, even within Europe, where the cycles are typically highly synchronised , there are still exceptions [8]- even on relatively small spatial scales [9]. This juxtaposition of evidence suggests further experimental research is needed to be done in order to understand the role of the global 3.5 year temperature cycle and its links to small rodent dynamics.

#### VI. EXTRA CLIMATE PLOTS

Below are the plots for all other climate variables over time.

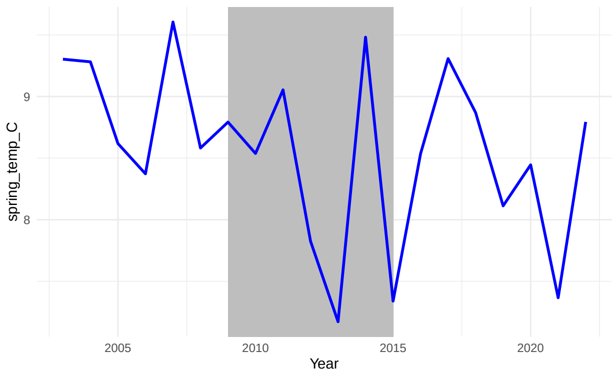


(a) Spring Temperatures in Glen Finglas.

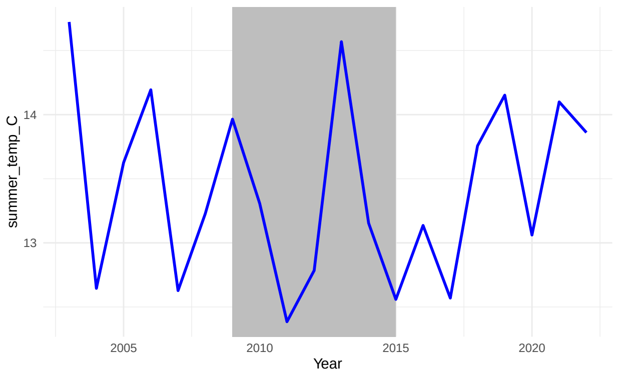


(b) Summer Temperatures in Glen Finglas.

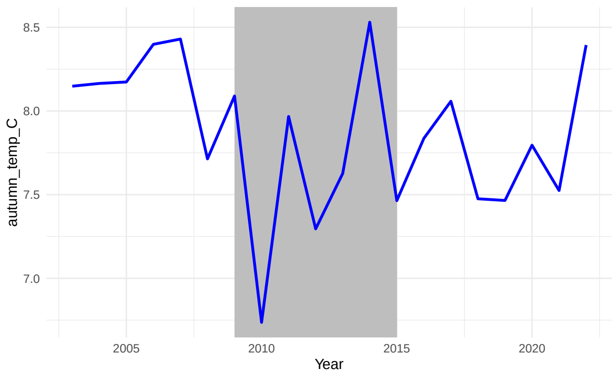


(c) Autumn Temperatures in Glen Finglas.

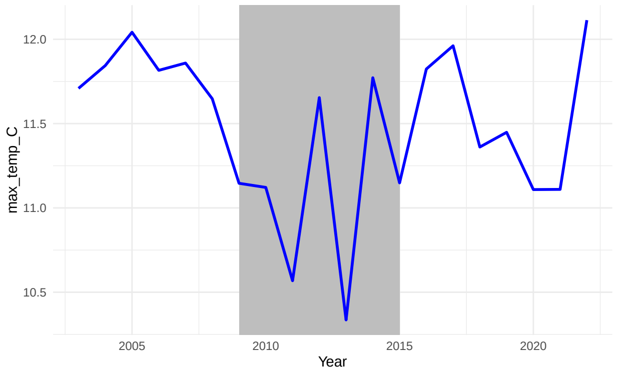


(d) Mean Maximum Daily Temperatures in Glen Finglas.

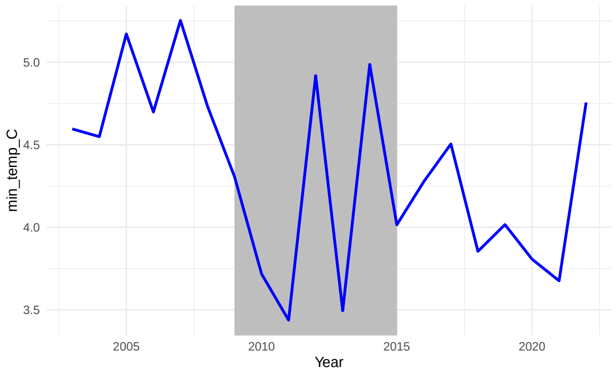


(e) Mean Minimum Daily Temperatures in Glen Finglas.

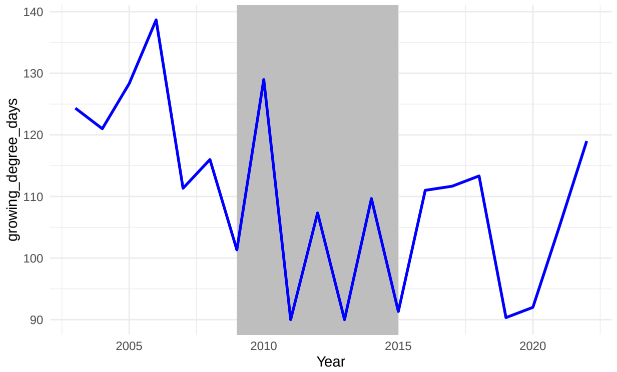


(f) Growing degree days in Glen Finglas.
